# Does stress make males? An experiment on the role of glucocorticoids in anuran sex reversal

**DOI:** 10.1101/2023.05.27.541969

**Authors:** Veronika Bókony, Csenge Kalina, Nikolett Ujhegyi, Zsanett Mikó, Kinga Katalin Lefler, Nóra Vili, Zoltán Gál, Caitlin R. Gabor, Orsolya Ivett Hoffmann

## Abstract

Environmentally sensitive sex determination may help organisms adapt to environmental change but also makes them vulnerable to anthropogenic stressors, with diverse consequences for population dynamics and evolution. The mechanisms translating environmental stimuli to sex are controversial: although several fish experiments supported the mediator role of glucocorticoid hormones, results on some reptiles challenged it. We tested this hypothesis in amphibians by investigating the effect of corticosterone on sex determination in agile frogs (*Rana dalmatina*). This species is liable to environmental sex reversal whereby genetic females develop into phenotypic males. After exposing tadpoles during sex determination to waterborne corticosterone, the proportion of genetic females with testes or ovotestes increased from 11% to up to 32% at 3 out of 4 concentrations. These differences were not statistically significant except for the group treated with 10 nM corticosterone, and there was no monotonous dose-effect relationship. These findings suggest that corticosterone is unlikely to mediate sex reversal in frogs. Unexpectedly, animals originating from urban habitats had higher sex-reversal and corticosterone-release rates, reduced body mass and development speed, and lower survival compared to individuals collected from woodland habitats. Thus, anthropogenic environments may affect both sex and fitness, and the underlying mechanisms may vary across ectothermic vertebrates.

## 1. Introduction

In many organisms, sex determination is influenced by the environment [1]. When there is also a genetic component of sex determination, for example in the form of sex chromosomes, its effects can be overridden by environmental influences such as thermal and chemical stressors, resulting in sex reversal: a mismatch between genetic sex and phenotypic sex [2]. Although environmentally sensitive sex determination (ESD) may have evolved as an adaptive sex-allocation strategy [3,4], in our era of rapid human-induced environmental changes it increases vulnerability to climate change, chemical pollution, the urban heat island effect, and other habitat alterations such as microclimatic changes due to invasive species [5–7]. The potential consequences are diverse, including the evolution of sex-determination systems and mating preferences, and skewed adult sex ratios [8–11] which can affect a wide range of social behaviours, life histories, reproductive systems, population viability and adaptive potential [7,12]. To understand and forecast these evolutionary-ecological outcomes, we need to find out what proximate mechanisms translate environmental stimuli into phenotypic sex.

Usually, the sex-determining environmental signals are stressful stimuli, like relatively extreme temperatures or pH, salinity, hypoxia, starvation, crowding, and altered light conditions [3,13–15]. Therefore, the glucocorticoid stress axis lends itself as a potential mediator of ESD. Glucocorticoid hormones are circulated at elevated concentrations in response to stressors and may affect the downstream pathways of sex differentiation, thereby affecting phenotypic sex [15]. This hypothesis has been supported by experimental evidence in various species of fish (Table 1). However, similar experiments in reptiles either did not support the role of glucocorticoids in sex determination or produced controversial results (Table 1). Instead, other molecular mediators of ESD have been suggested for both taxa [14,16–18]. To date, it remains unresolved whether ESD is regulated by a universal mechanism or by different signal-transducer systems in different clades.

**Table 1.** Published effects of embryonic or larval treatment by glucocorticoids (GC; cortisol in fish, corticosterone in reptiles and amphibians) on phenotypic sex ratios in ectothermic vertebrates.

| Class | Species | Sex chromosomes | Temperature effect on sex<sup>*</sup> | GC effect on sex<sup>*</sup> | Notes | Reference<sup>‡</sup> |
| --- | --- | --- | --- | --- | --- | --- |
| Fish | *Odontesthes bonariensis* | XX/XY | low F, high M | M | references on sex-determination system: [52,53] | [45] |
| Fish | *Oncorhynchus mykiss* | XX/XY | none or family-specific | M | references on sex-determination system: [54,55] | [56] |
| Fish | *Oryzias latipes* | XX/XY | high M | M | GC might act through oxidative stress [47] | [33] |
| Fish | *Paralichthys lethostigma* | XX/XY | low and high M | M | blue tank colour increases GC and the ratio of males | [57] |
| Fish | *Paralichthys olivaceus* | XX/XY | high M | M |  | [58] |
| Reptiles | *Amphibolurus muricatus* | none identified | low and high F | F | GC effect may be due to sex-specific mortality | [59] |
| Reptiles | *Bassiana duperreyi* | XX/XY | low M | M | GC effect might be due to sex-specific mortality | [59] |
| Reptiles | *Ctenophorus fordi* | ZW/ZZ | none found (but small sample size) | no |  | [60] |
| Reptiles | *Pogona vitticeps* | ZW/ZZ | high F | no |  | [44] |
| Reptiles | *Trachemys scripta* | none identified | low M, high F | no | GC was tested at male-producing temperature only | [61] |
| Amphibians | *Rana dalmatina* | XX/XY | high M | no | sex-reversal rate (M) increased from 11% to 32% at 1 out of 4 GC concentrations | This study |

\**Low* and *high* refer to environmental temperatures; M and F signify resulting sex-ratio shifts towards males and females, respectively.
‡For references on sex-determination systems in reptiles and amphibians, see the HerpSexDet database [62]. For fish, information on sex- determination systems can be found in the references cited here for the GC effect on sex, unless stated otherwise in the Notes column.

Studying this question in amphibians may take us one step closer to the answer. Amphibians resemble fishes in several relevant aspects: they are anamniotes and their sex determination usually takes place in freely moving larvae that are directly exposed to the environment, rather than developing in shelled eggs within a nest or in the mother’s body. Both fish and amphibians typically respond to environmental stress with an over-production of phenotypic males (female-to-male sex reversal or type-1b temperature-dependent sex determination) as opposed to the more variable reaction norms among reptiles [2]. On the other hand, amphibians resemble reptiles in that they are gonochoristic, meaning that phenotypic sex does not change throughout individual lifetime, since the ontogeny of reproductive and urinary tracts are intertwined, as opposed to fish that are capable of sequential hermaphroditism [19]. Also, the main glucocorticoid hormone is corticosterone in amphibians and reptiles but cortisol in fish. Finally, amphibians differ from both reptiles and fish in that genetic sex determination seems to prevail in amphibians and the environment affects their sex only outside the “normal” range of conditions, whereas in many reptiles and fishes the environmental effect is strong under natural environmental conditions and the genetic component is weak or apparently absent [2].

To our knowledge, no study has yet been published on the effect of biologically active glucocorticoids on sex determination in amphibians [20]. Therefore, here we examined the role of corticosterone (CORT), the main glucocorticoid hormone of amphibians, in sex reversal in the agile frog (*Rana dalmatina*). This anuran species has male-heterogametic sex determination (XX/XY sex chromosomes) system [21], but larval heat stress causes female-to-male sex reversal [22,23], and ca. 20% of phenotypic males in natural populations are genetically females [6]. We treated tadpoles with CORT during their most sensitive period of sex determination to test if this hormone triggers sex reversal.

## 2. Methods

### (a) Experimental protocol

We collected freshly spawned agile frog eggs from each of three egg masses from each of four sites (populations) in Hungary on 7 April 2021. We sampled four populations to facilitate the generalizability of results: two sites are in the cities of Budapest (Erzsébet-ér: N 47.429196, E 19.132845) and Göd (Feneketlen-tó: N 47.684865, E 19.129855), respectively; two sites are in woodlands (Bajdázó-tó: N 47.903382, E 18.978482; Ilona-tó: N 47.713432, E 19.040242). We reared the embryos in our laboratory, each of the twelve sibling groups kept in a separate container with ca. 1 cm high reconstituted soft water (RSW; 48 mg NaHCO_3_, 30 mg CaSO_4_ × 2 H_2_O, 61 mg MgSO_4_ × 7 H_2_O, 2 mg KCl added to 1 L reverse-osmosis filtered, UV-sterilized, aerated tap water). Throughout the study, air temperature in the laboratory was 19°C and we regularly adjusted the photoperiod to mimic the outdoors dark-light cycles.

When the animals reached developmental stage 25 [24], we haphazardly selected 25 tadpoles from each sibling group; the remaining tadpoles were released at their ponds of origin. We placed each of the 300 experimental tadpoles into a white plastic container filled with 1 L RSW, arranged in a randomized block design to ensure that all five treatments and all four populations were homogeneously distributed across the shelves in the laboratory. We randomly assigned each tadpole to a treatment group (see below) in a stratified way such that there were five tadpoles from each sibling group (15 animals per population) in each group, yielding 60 individuals in total for each treatment. Twice per week throughout the experiment, we changed the rearing water and fed the tadpoles *ad libitum* with slightly boiled chopped spinach.

The treatment period was the third week (14^th^ – 20^th^ days) of tadpole development, which corresponds to the most sensitive period of sex determination in agile frogs [22]. During this period, we changed the rearing water every other day, and each tadpole received chemically treated RSW according to its assigned treatment. The control group received 0.1 mL/L ethanol into their rearing water; this ethanol concentration is much lower than those observed to harm anuran embryos or tadpoles [25–27]. The same amount of ethanol was also present as solvent in the rearing water of the remaining four treatment groups, which received 0.01, 10, 100, or 1000 nM CORT, respectively. We chose these concentrations for the following reasons. The lowest concentration represents the ecologically relevant range of CORT reported from surface waters that serve as tadpole-development habitat of many amphibians [28,29]. The two medium concentrations represent the biologically active range for acute stress, based on the previous findings that waterborne CORT treatments of these magnitudes resulted in endogenous CORT levels similar to those measured after exposure to natural stressors such as starvation, crowding, and predation risk [30–32]. The highest concentration is a pharmacological dose that we chose for comparability with earlier experiments on cortisol in fish [33,34] and corticosteroid metabolites in amphibians [20]. We renewed each treatment every 48 hours [35], and we ended all treatments after six days by changing the rearing water of all tadpoles to clean RSW.

### (b) Corticosterone measurement

Immediately after the water change that ended all treatments, we sampled each tadpole for CORT release rate using a non-invasive waterborne hormone sampling method, which provides an integrated CORT measurement that is repeatable within individuals and correlates with plasma concentrations [31,36,37]. We transferred the tadpoles from their rearing containers with single-use mesh nets to individual 0.5 L plastic cups containing 100 ml RSW. After each tadpole spent one hour in the cup, we measured their body mass (± 0.1 mg), we poured the water through filter paper (coffee filters equivalent to grade 4 filter paper; to remove faeces) into individual containers and stored the water samples at -20 °C until analysis. We had 287 samples, because 7 tadpoles died before sampling and 6 samples were accidentally lost during sampling.

We extracted CORT from the water-borne hormone samples following established protocol [29]. Briefly, we used C18 SPE columns (Sep-Pak Vac, 3cc/500mg, Waters Inc.) to extract the CORT and then we froze the samples at -20 °C. Later we eluted samples with 4 ml methanol and dried them with nitrogen gas. Samples were analysed using a competitive enzyme-linked immunosorbent assay (ELISA) kit (№ 501320, Cayman Chemical Inc.) in duplicate. We resuspended each sample in 25 μl of 95 % ethanol followed by 475 μl ELISA buffer (total 500 μl). We measured sample absorbance on a fluorescent plate reader (800XS; Biotek Instruments Inc.) at 405 nm. Inter-assay coefficient of variation (CV) of the concentration standards was 11.7 %. Intra-assay CV of the tadpole samples ranged from 8.4 to 21 % across 14 plates (median: 9.4, mean: 11.76%). All but 2 plates had <15 % CV; 9 had ≤10 % CV. Samples with high intra-assay CV were re-run and we used the measurement with the lowest CV for each tadpole. Four out of the 287 tadpoles were excluded from the statistical analyses because their CORT concentration was outside the range of the calibration curve. We calculated the hourly CORT release rate by multiplying CORT (pg/ml) by the resuspension volume (0.5 ml) and dividing by the mass of each resources tadpole (pg/mg/h).

### (c) Sex identification

To diagnose phenotypic sex, we raised the animals to metamorphosis and two months thereafter. When an individual reached developmental stage 42 (start of metamorphosis: the appearance of forelimbs), we recorded the date and measured metamorphic body mass (± 0.1 mg). We slightly tilted the container and decreased the water level to 0.1 L to allow the animal to leave the water. Upon completion of metamorphosis (developmental stage 46: disappearance of the tail), we moved the froglet into a clean rearing box that contained wet paper towels as substrate and a piece of egg carton as shelter, which we changed every other week. The froglets were fed *ad libitum* with springtails and small (2-3 mm) crickets, sprinkled with a 3:1 mixture of Reptiland 76280 (Trixie Heimtierbedarf GmbH & Co. KG, Tarp, Germany) and Promotor 43 (Laboratorios Calier S.A., Barcelona, Spain) containing vitamins, minerals and amino acids.

We dissected the froglets for phenotypic sexing ca. two months after metamorphosis (124-175 days after starting the experiment at stage 25); at this age the gonads are well differentiated in agile frogs [23]. We measured the froglets’ body mass (± 0.01 g) and euthanized them using a one-hour water bath containing 6.6 g/L MS-222 buffered to neutral pH with the same amount of Na_2_HPO_4_. Because the digestive tract sometimes contained food remains, we measured its mass to subtract it from body mass for the statistical analyses. We examined and photographed the gonads under an APOMIC SHD200 digital microscope to record whether the individual had testes or ovaries. We stored the gonads in neutral-buffered 10% formalin until histological analysis, which we performed by routine methods as described in our earlier papers [6,23]. Histological section preparation was successful for 193 out of 203 dissected individuals. Using the information from both dissection and histology, we categorized phenotypic sex as male, female, or intersex (individuals with ovotestes, i.e. multiple oogonia in testicular tissue).

For genetic sexing, we removed the hind feet of each froglet after euthanasia, using scissors and forceps that we sterilized for each individual by dipping into ethanol and flaming. We stored the tissue samples in 96% ethanol until DNA extraction, which we performed using E.Z.N.A. Tissue DNA Kit following the manufacturer’s protocol, except that digestion time was 2 hours. For identifying genetic sex, we used the method of Nemesházi et al. [6]. Briefly, we tested all froglets for sex marker Rds3 (≥ 95% sex linkage; primers: Rds3-HRM-F and Rds3-HRM-R) using high-resolution melting (HRM), and we accepted an individual to be concordant male or concordant female if its Rds3 genotype was in accordance with its phenotypic sex. Those individuals that appeared sex-reversed by Rds3 were also tested for sex marker Rds1 (≥ 89% sex linkage; primers: Rds1-F, Rds1-R and Rds1-Y-R) using PCR and were accepted to be sex-reversed only if both markers confirmed sex reversal. All dissected individuals were successfully genotyped, and the two markers gave the same genetic sex in each case where Rds1 was tested.

Over the experiment, mortality was unexpectedly high: 81 individuals (27%) died for unknown reasons (3, 4, and 74 deaths before, during, and after treatment, respectively).

Our sample size was further reduced by 16 urban tadpoles that did not start metamorphosis by the time of dissection; these animals could not be phenotypically sexed because their gonads were not developed enough. Additionally, one dissected froglet (an urban individual in the highest-concentration CORT treatment group) had under-developed gonads and could not be unambiguously sexed phenotypically (it was genetically male). Therefore, we had sufficient data from 202 individuals for the analysis of sex reversal, 119 of which were genetically female.

### (d) Statistical analyses

We analysed the data using the R computing language [38] with packages ‘geepack, ‘coxme’, and ‘emmeans’. To evaluate the effects of CORT treatment on the tadpoles’ CORT release rate (log10-transformed for the analyses) and body mass at the three investigated life stages (end of treatment period, metamorphosis, and dissection), we ran a Generalized Estimating Equations (GEE) model for each of the four dependent variables. We used treatment and site of origin as predictors; additionally, we included age at dissection (from the completion of metamorphosis) in the model of body mass at dissection. To analyse the effect of CORT treatment on sex-reversal rate, we used the subset of genetically female individuals (XX genotypes), and we applied a binomial GEE model on the presence of sex reversal (male or intersex phenotype as opposed to female), using treatment and site of origin as predictors. In all GEE models, we allowed for the non-independence of siblings by using the ‘exchangeable correlation’ (or ‘compound symmetry’) association structure [39].

To test the effect of CORT treatment on survival over the experiment and time to metamorphosis, we ran a mixed-effects Cox’s proportional hazards model for each of these two dependent variables. Individuals that were euthanized at the end of experiment, or those that did not start metamorphosis, respectively, were used as censored observations. Both models included treatment and site of origin as fixed factors, and sibling group as random factor.

We used diagnostic residual plots to ascertain that the data fit the statistical assumptions of each analysis. From each model, we tested the difference of each CORT treatment group from the control group by estimating linear contrasts of marginal means, and for these multiple comparisons we corrected the P-values with the false discovery rate (FDR) method [40]. For comparisons between sites of origin, we calculated a single contrast between the two urban sites and the two woodland sites from each model. We did this post-hoc comparison because we noted differences between the urban and woodland populations during the experiment, although testing the effects of site or habitat was not an *a-priori* goal of the study. The R script of our statistical analyses, annotated with detailed comments, are provided as supplementary material.

## 3. Results

At the end of treatment, CORT release rate was ca. 3 times higher in the group treated with the highest CORT concentration compared to the control group, but there was no significant difference in the three lower-concentration treatment groups (table 2, figure 1). However, body mass at the end of treatment was significantly reduced by all CORT concentrations (marginally so by the lowest concentration), and this effect increased with concentration (table 3, figure 1). There were no treatment effects on body mass at metamorphosis or at dissection (table 3). CORT treatment had no significant effect on survival (65-76.7% per group survived) or time to metamorphosis (table 2, figures S1-S2). The frequency of sex reversal was 11% in the control group and varied between 11% and 32% in the CORT-treatment groups; these differences did not follow a concentration-dependent pattern (figure 2). The effect of CORT treatment on sex-reversal rate was statistically non-significant, except for the group treated with 10 nM CORT (table 2).

**Table 2.** Effects of CORT treatment and habitat of origin on CORT release rate, survival over the experiment, time to metamorphosis, and female-to-male sex-reversal rate. For each dependent variable, pairwise comparisons are given as ratios, i.e. proportional differences ± standard error calculated from marginal means estimated from the model. P-values are corrected for false discovery rate; P > 0.05 values are highlighted in bold type.

| Comparisons | CORT release rate<sup>*</sup> |  | Survival<sup>‡</sup> |  | Metamorphosis<sup>‡</sup> |  | Sex reversal<sup>†</sup> |  |
| --- | --- | --- | --- | --- | --- | --- | --- | --- |
|  | Ratio | P | Ratio | P | Ratio | P | Ratio | P |
| 0.01 nM / control | $0.85 \pm 0.18$ | 0.521 | $1.38 \pm 0.47$ | 0.918 | $0.90 \pm 0.19$ | 0.627 | $2.02 \pm 1.67$ | 0.395 |
| 10 nM / control | $0.89 \pm 0.15$ | 0.521 | $0.96 \pm 0.35$ | 0.918 | $0.89 \pm 0.18$ | 0.627 | $3.87 \pm 2.06$ | **0.043** |
| 100 nM / control | $1.13 \pm 0.21$ | 0.521 | $1.04 \pm 0.37$ | 0.918 | $0.69 \pm 0.15$ | 0.159 | $0.80 \pm 0.17$ | 0.394 |
| 1000 nM / control | $2.99 \pm 0.57$ | **<0.001** | $0.90 \pm 0.33$ | 0.918 | $0.64 \pm 0.13$ | 0.115 | $1.69 \pm 0.76$ | 0.394 |
| Urban / woodland | $2.17 \pm 0.28$ | **<0.001** | $7.71 \pm 2.74$ | **< 0.001** | $0.11 \pm 0.03$ | **< 0.001** | $6.48 \pm 5.43$ | **0.026** |

\* Ratios of group means are given from Generalized Estimating Equations model; N = 287.
‡ Hazard ratios are given from Cox's proportional hazards models; N = 300.
† Odds ratios are given from Generalized Estimating Equations model; N = 119.

**Table 3.** Effects of CORT treatment and habitat of origin on body mass at the end of treatment, at metamorphosis, and at dissection. Pairwise comparisons (differences between group means ± standard error, SE) are taken from three Generalized Estimating Equations models. P-values are corrected for false discovery rate; P > 0.05 values are highlighted in bold type. Sample sizes are 293, 243, and 202, respectively.

| Life stage | Comparisons | Difference ± SE | P |
| --- | --- | --- | --- |
| End of treatment | 0.01 nM - control | -18.1 ± 10.0 | 0.071 |
| | 10 nM - control | -21.2 ± 8.8 | **0.021** |
| | 100 nM - control | -56.0 ± 15.5 | **0.001** |
| | 1000 nM - control | -63.9 ± 11.6 | **<0.001** |
| | Urban - woodland | -188.0 ± 13.8 | **<0.001** |
| Metamorphosis | 0.01 nM - control | -12.4 ± 11.1 | 0.515 |
| | 10 nM - control | -7.0 ± 13.1 | 0.594 |
| | 100 nM - control | -8.1 ± 9.4 | 0.515 |
| | 1000 nM - control | -20.8 ± 16.6 | 0.515 |
| | Urban - woodland | -89.3 ± 22.7 | **<0.001** |
| Dissection | 0.01 nM - control | -61.8 ± 43.0 | 0.602 |
| | 10 nM - control | -12.8 ± 44.5 | 0.776 |
| | 100 nM - control | 8.5 ± 30.0 | 0.776 |
| | 1000 nM - control | 23.7 ± 43.4 | 0.776 |
| | Urban - woodland | 7.9 ± 33.9 | 0.814 |

**Figure 1.**
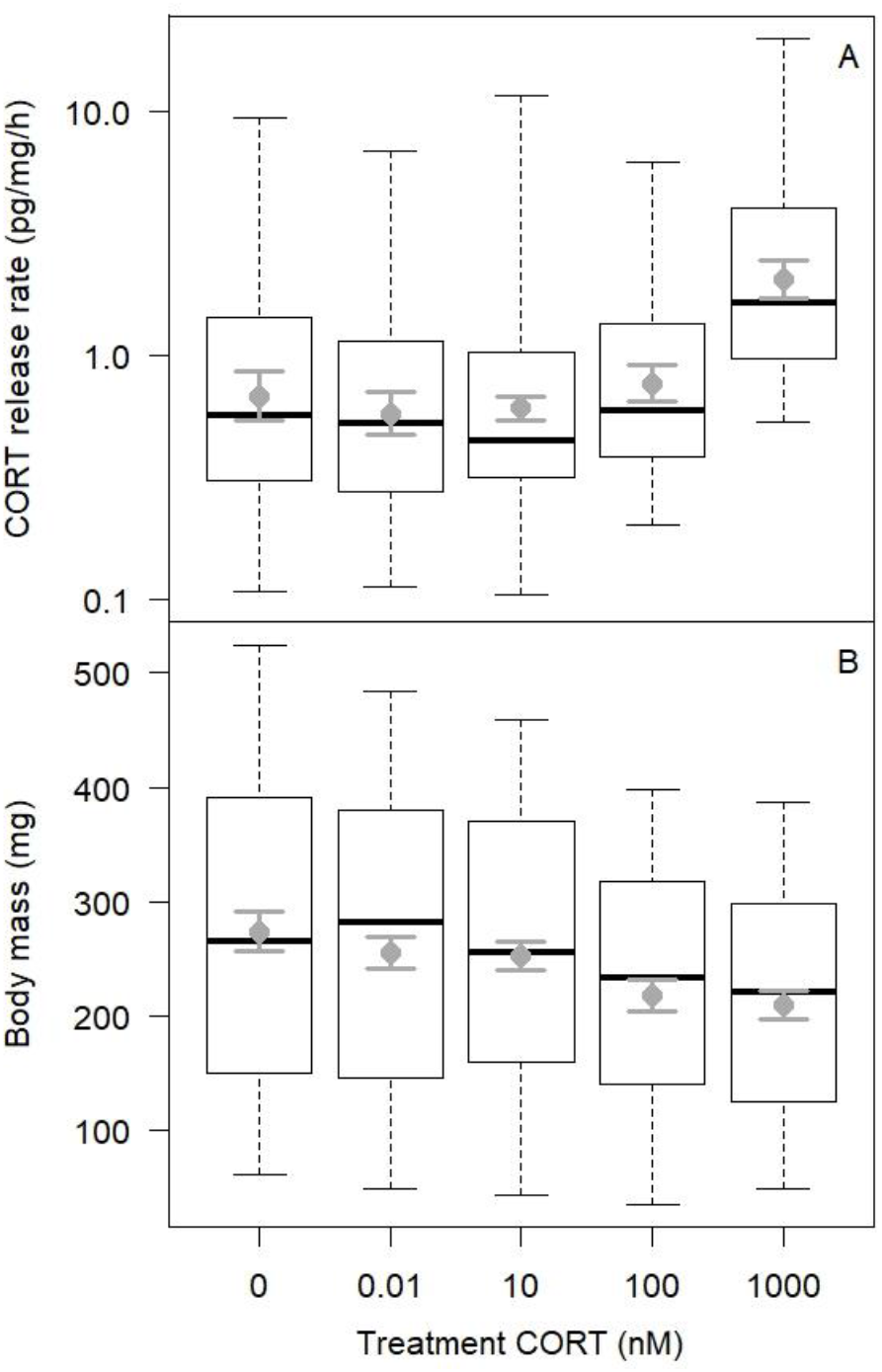
Corticosterone (CORT) release rate and body mass of tadpoles at the end of CORT treatment in each group. In each boxplot, the thick horizontal line, box, and whiskers represent the median, interquartile range, and data range, respectively. Dots with error bars show the model-estimated means with 84% confidence intervals; non-overlapping error bars correspond to significant differences at α=0.05 after correction for false discovery rate.

**Figure 2.**
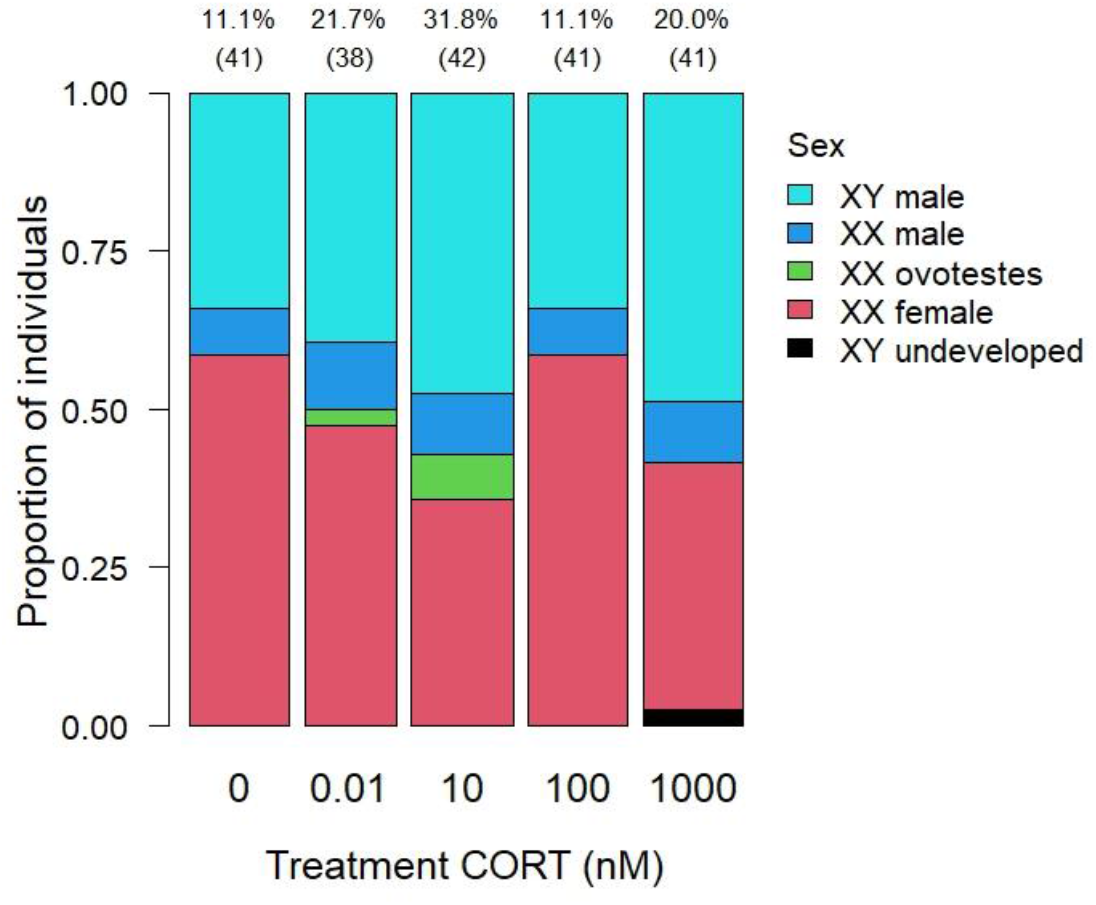
Combinations of genetic and phenotypic sex in the CORT treatment groups. Percentage of sex reversal (individuals with testes or ovotestes within XX genotypes) is given above each bar, along with sample size (total number of sexed individuals) in parentheses.

Almost all endpoints measured in the experiment were significantly affected by the type of habitat the animals originated from. Specifically, the probability of female-to-male sex reversal was higher in froglets originating from the two urban sites compared to their counterparts from the two woodlands (table 2, figure 3). Also, individuals from urban sites had higher CORT release rates, smaller body mass at the end of treatment and at metamorphosis (but not at dissection), longer time to metamorphosis, and higher mortality compared to the woodland individuals (tables 2-3, figure 3, figures S3-S4). These latter differences were not due to the higher proportion of sex-reversed individuals in urban populations, because there was no significant difference between sex-reversed and concordant individuals in CORT release rate, body mass, and time to metamorphosis (table S1, figures S5-S8).

**Figure 3.**
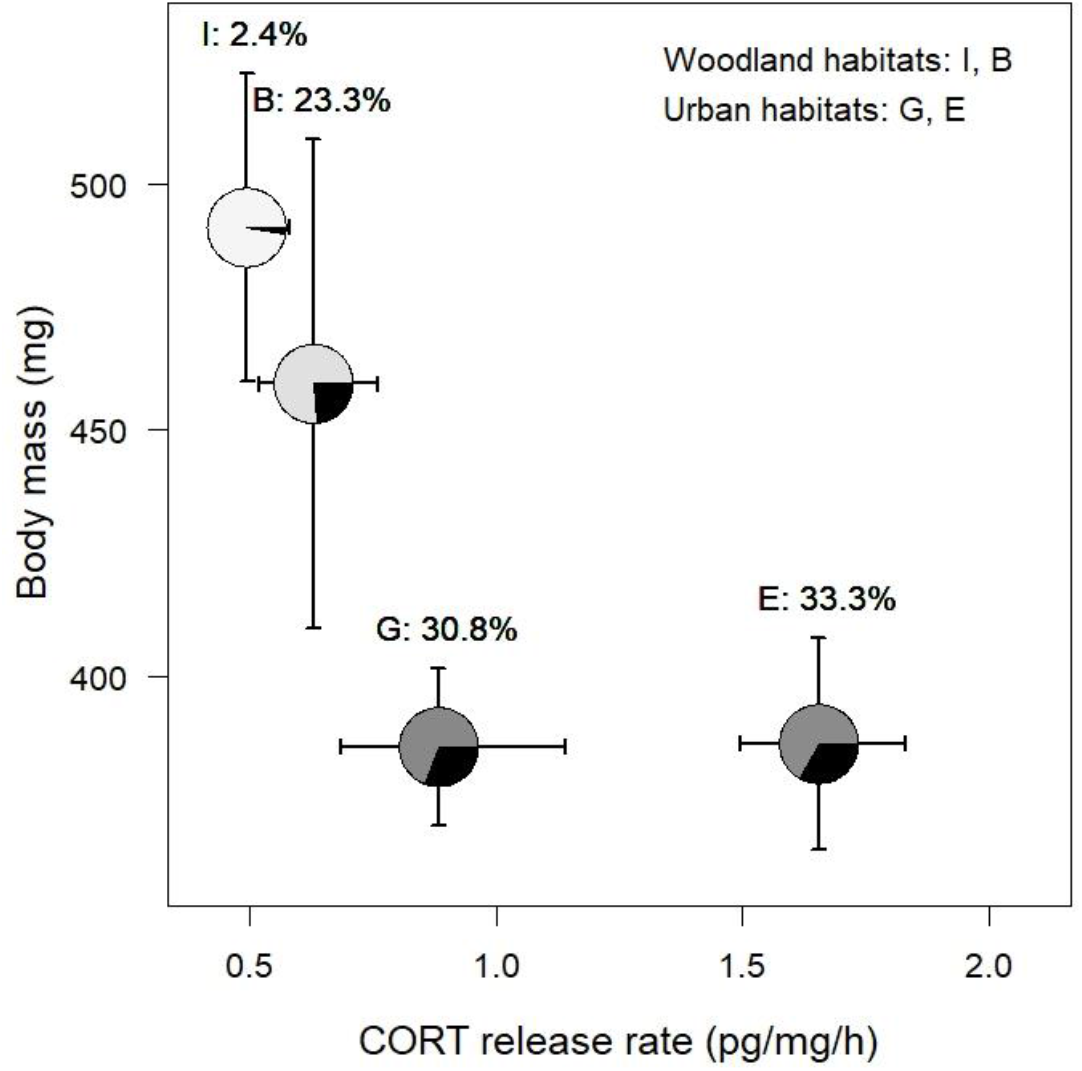
Corticosterone (CORT) release rate at the end of treatment and body mass at metamorphosis by the site of origin. In each pie chart, the size of the black section indicates the rate of sex reversal (percentage of XX individuals with testes or ovotestes; written above each pie chart), and the grey colour indicates survival over the experiment, with darker grey representing higher mortality (4, 12, 46.7, and 45.3 %, respectively, in populations I, B, G, and E). Error bars around pie charts show the 84% confidence interval of each population’s mean; non-overlapping error bars correspond to significant differences at α=0.05 after correction for false discovery rate. Note that error bars for CORT are asymmetrical because they are back-transformed from the log10-scale.

## 4. Discussion

The main result of our experiment is that we found no convincing effect of CORT treatment on sex-reversal rate in agile frogs. Although the proportion of genetic females with testes or ovotestes was ca. 2-3 times higher in 3 out of 4 treated groups compared to the control group, these differences were not statistically significant except for the group treated with 10 nM CORT and did not align to a monotonous dose-effect relationship. This suggests that CORT is not likely to be the mediator of female-to-male sex reversal in this anuran species.

Before discussing the implications of this finding, we should consider the alternatives. First, it is possible that CORT treatment was ineffective in elevating the internal CORT levels of tadpoles. We cannot rule out this option because we did not directly measure circulating CORT concentrations in our animals, although the CORT release rates that we measured with our waterborne method correlate with internal CORT levels in many other amphibian species [41]. In any case, CORT release rate was significantly elevated in the group treated with 1000 nM CORT; thus, we should have observed increased sex-reversal rate in this group if CORT had affected sex determination. Furthermore, tadpole body mass at the end of treatment was reduced in all CORT-treated groups in a monotonously concentration-dependent manner, which indirectly supports the effectiveness of these treatments because of the growth-inhibiting effect of CORT in young tadpoles [32,42,43]. The finding that CORT levels during sex determination did not reflect CORT treatments in our experiment is similar to discrepancies reported in previous studies, which the authors attributed to methodological difficulties with CORT measurement, e.g. due to rapid clearance [44,45].

Second, our findings on sex reversal may be complicated by the unexpectedly high mortality after the CORT treatments. First, the ca. 30% decrease in overall sample size reduced our statistical power and the precision of our inferences. Second, it could have biased our data if sex reversal had affected mortality. The latter possibility is unlikely to account for our results, however, because mortality was independent of CORT treatment. Furthermore, we found no effect of genetic sex on mortality in an earlier study of agile frog tadpoles and juveniles [46]. Altogether, we conclude that mortality and CORT-methodological issues might have added some noise to our data but, had there been a strong treatment effect, such an effect would have been unlikely to get masked by noise. Compared to the strong effect of a physiological stressor, heat exposure, which caused >90 % sex reversal in previous experiments [22,23], the rates of sex reversal in the present study (11-32 %) were low. Therefore, the variation among treatment groups is probably attributable to random deaths combined with the low but steady “baseline” sex-reversal propensity in agile frogs [22,46] rather than to a genuine effect of CORT treatment.

Arriving at the conclusion that CORT did not drive sex reversal in agile frogs, we can speculate that the proximate mechanism of ESD in amphibians may be something else than the glucocorticoid pathway supported in fishes. Proposed alternative mechanisms (none of which are mutually exclusive) are the cellular calcium and redox regulation [16,47], epigenetic processes [18,48], thyroid hormones [15,20,49], and the energy limitation hypothesis [14]. The latter proposes that nutritional stress *via* lipid and carbohydrate metabolism pathways serves as signal transducer between environment and sex. At first glance, our results might support this idea, because the incidence of sex reversal correlated negatively with body mass across the four populations (Fig. 3). However, this relationship was not detectable within populations at the level of individuals (table S1, figures S5-S8), suggesting that the among-population differences in sex-reversal rate and in the other phenotypic traits were not causally related but rather caused by some other factor. This potential other factor was strongly linked with the habitat type of the populations from which our experimental animals originated. Specifically, individuals from urban sites had reduced performance in terms of survival, body mass, and development speed, similarly to earlier findings on another anuran species [50]. These differences were accompanied by higher sex-reversal rates in the frogs from urban sites in the present study, mirroring the sex-reversal rates observed earlier in the free-living adults at the same sites [6]. One potential explanation to these findings is that urbanization may affect both offspring vigour and the propensity for sex reversal independently from each other, for example *via* epigenetic effects of chemical pollutants [6,50,51]. Alternatively, sex reversal may be causally related to reductions in fitness-related traits [23,46], and the environmental stressors associated with urbanization may trigger this cause-effect cascade [51]. To tease apart these possibilities, we urge further studies designed specifically to test the inter-relationships among environmental stress (especially by human-induced environmental changes), sexual fate, and individual fitness in ESD species.

To sum up, our study did not support the role of glucocorticoids in anuran sex reversal, tentatively placing amphibians together with reptiles and apart from fishes in terms of the proximate drivers of environmental sex determination. However, considering the very small number of species studied from each taxonomic class (Table 1), it remains an open question if the heterogeneity observed so far has anything to do with phylogeny. For example, all species in which a male-biasing effect of glucocorticoid treatment was reported have male-heterogametic sex-chromosome systems and all but one of them have ESD where extreme temperatures produce males (Table 1). This pattern is contradicted by our present results, however (Table 1), highlighting the unreliability of generalizing from a few results. Clearly, more studies are needed on a more diverse array of species to understand how environmental stress is translated into phenotypic sex across ectothermic vertebrates.

## Supporting information

Supplementary table & figures, data, and R script.

## Ethics

This study was approved by the Ethics Committee of the Plant Protection Institute and licensed by the Environment Protection and Nature Conservation Department of the Pest County Bureau of the Hungarian Government (PE-06/KTF/8060-1,2,3/2018; PE/EA/295-7/2018).

## Data accessibility

The dataset and R script supporting this article have been uploaded as part of the supplementary material.

## Authors’ contributions

V.B.: conceptualization, data curation, formal analysis, funding acquisition, investigation, project administration, supervision, visualization, writing – original draft and review & editing. C.K.: data curation, investigation. U.N.: data curation, investigation. Z.M.: investigation. K.K.L.: investigation. N.V.: investigation. Z.G.: investigation. C.R.G.: funding acquisition, supervision, writing – review & editing. O.I.H.: writing – original draft and writing – review & editing.

## Funding

The study was supported by the National Research, Development and Innovation Office of Hungary (NKFIH K-135016, ÚNKP-21-5, and 2019-2.1.11-TÉT-2019-00026 to VB, NKFIH PD-134241 to ZM, NKFIH FK-124708 to OIH) and the János Bolyai Scholarship of the Hungarian Academy of Sciences to VB and OIH.

## Acknowledgements

We thank all members of the Evolutionary Ecology Research Group for their help during the experiment, and Saeid Panahi Hassan Barough for help with hormone extractions and plating.

## Notes

### Competing Interest Statement

The authors have declared no competing interest.

