## Supplementary table & figures, data, and R script. for "Does stress make males? An experiment on the role of glucocorticoids in anuran sex reversal": Bokony_Annotated R script CORTsex.pdf

**Annotated R script for the analyses in the paper "*Does stress make males? An experiment on the role of glucocorticoids in anuran sex reversal*" by Veronika Bókony et al.**

Contact:

```
# The codes below were run with R version 4.2.1

# Set working directory and read the data file:
setwd("D:/Your_working_folder")
i=read.csv("Data_Bokonyetal_cort_sex.csv", header=T, sep=";", stringsAsFactors=TRUE)

# Re-order factor levels for "site" and order the dataset by this factor:
i$site =factor(i$population, levels=c("I","B","G","E"))
i=i[order(i$sibgroup),]

# Assign labels to factor levels:
i$treatment =factor(i$treatment, labels=c("0 nM", "0.01 nM","10 nM","100 nM","1000
    nM"))
i$sex.rev.3 =factor(i$sex.rev.3, labels=c("XY male", "XX male", "XX female"))
i$sex.rev.hist = factor(i$sex.rev.hist, levels=c("undeveloped", "XX_female",
    "XX_ovotestis", "XX_male", "XY_male"), labels=c("XY undeveloped", "XX female", "XX
    ovotestes", "XX male", "XY male"))

# STATISTICAL ANALYSES:
# Load libraries:
library(emmeans)
library(coxme)
library(geepack)
library(DHARMa)
library(survminer)
library(lattice)
library(plotrix)
```

### **# Tadpole CORT release rate**

```
# Diagnostics for Gaussian model:
plot(lm(log10(cort.rate) ~ treatment + site, data=i))
# Generalized Estimating Equations (GEE) model with Gaussian distribution, taking into
   account the non-independence of siblings by using the 'exchangeable correlation'
   structure. The benefit of this approach is that it offers greater power for
   estimating population averages when the levels of the random factor are nested
   within the levels of the fixed factor (in our case, sibling groups within sites of
   origin) as opposed to more widely used mixed-effects models. The drawback is that
   GEE provides no estimates for random intercepts; however, we did not aim to do so as
   we were not interested in the specific sibgroup effects.
gee2= geeglm(log10(cort.rate) ~ treatment + site, family=gaussian, id=sibgroup,
   corstr="exchangeable", data=i[!is.na(i$cort.rate),])
# Treatment effects:
(e.gee2.treat=emmeans(gee2, trt.vs.ctrl1~treatment, adjust="fdr", type="response"))
# Site effects:
 (e.gee2=emmeans(gee2, pairwise~site, adjust="fdr", type="response"))
# Difference between urban and woodland habitats:
contrast(e.gee2$emmeans, list(urb_nat=c(-0.5,-0.5,0.5,0.5)))
# Do sex-reversed and concordant individuals differ?
```

```
gee2.sex= geeglm(log10(cort.rate) ~ sex.rev.2, family=gaussian, id=sibgroup,
   corstr="exchangeable", data=i[!is.na(i$cort.rate) & !is.na(i$sex.rev.2),])
emmeans( gee2.sex, pairwise~sex.rev.2, type="response")$contrasts
# Figure S5:
with(i, bwplot(cort.rate ~ sex.rev.3|site, data=i, layout=c(4,1), ylab="CORT release
   rate (pg/mg/h)"))
```

**# Body mass**

```
# Plotting body mass at the 3 life stages by sex and site:
# Creating a “long” dataset from the “wide” dataset:
ii = reshape(i, direction = "long", varying = list(c("mass.may12", "mass.meta",
   "mass.diss.net")), v.names = "mass", idvar = c("ID", "treatment", "population",
   "sibgroup"), timevar = "time", times = 1:3)
ii$time = factor(ii$time, labels = c("post-treatment", "metamorphosis", "dissection"))
# Figure S6:
with(i, bwplot(mass ~sex.rev.3| site * time, data=ii, ylab="Body mass (mg)"))
```

**# Body mass at end of treatment**

```
# Diagnostics for Gaussian model:
plot(lm(mass.may12 ~ treatment + site, data=i))
# GEE model with Gaussian distribution, taking into account the non-independence of
   siblings by using the ‘exchangeable correlation’ structure:
gee3= geeglm(mass.may12 ~ treatment + site, family=gaussian, id=sibgroup,
   corstr="exchangeable", data=i[!is.na(i$mass.may12),])
# Treatment effects:
(e.gee3.treat=emmeans(gee3, trt.vs.ctrl1~treatment, adjust="fdr", type="response"))
# Site effects:
(e.gee3=emmeans(gee3, pairwise~site, adjust="fdr", type="response"))
# Difference between urban and woodland habitats:
contrast(e.gee3$emmeans, list(urb_nat=c(-0.5,-0.5,0.5,0.5)))
# Do sex-reversed and concordant individuals differ?
gee3.sex= geeglm(mass.may12 ~ sex.rev.2, family=gaussian, id=sibgroup,
   corstr="exchangeable", data=i[!is.na(i$mass.may12) & !is.na(i$sex.rev.2),])
emmeans(gee3.sex, pairwise~sex.rev.2, type="response")$contrasts
```

**# Body mass at metamorphosis**

```
# Diagnostics for Gaussian model:
plot(lm(mass.meta ~ treatment + site, data=i))
# GEE model with Gaussian distribution, taking into account the non-independence of
   siblings by using the ‘exchangeable correlation’ structure:
gee4= geeglm(mass.meta ~ treatment + site, family=gaussian, id=sibgroup,
   corstr="exchangeable", data=i[!is.na(i$mass.meta),])
# Treatment effects:
(e.gee4.treat=emmeans(gee4, trt.vs.ctrl1~treatment, adjust="fdr", type="response"))
# Site effects:
(e.gee4=emmeans(gee4, pairwise~site, adjust="fdr", type="response"))
# Difference between urban and woodland habitats:
contrast(e.gee4$emmeans, list(urb_nat=c(-0.5,-0.5,0.5,0.5)))
# Do sex-reversed and concordant individuals differ?
gee4.sex= geeglm(mass.meta ~ sex.rev.2, family=gaussian, id=sibgroup,
   corstr="exchangeable", data=i[!is.na(i$mass.meta) & !is.na(i$sex.rev.2),])
```

```
emmeans(gee4.sex, pairwise~sex.rev.2, type="response")$contrasts
```

**# Body mass at dissection**

```
# Centering age at dissection to its mean:
i$dissect.age.z = scale(i$dissect.age, center = TRUE, scale = FALSE)
# Diagnostics for Gaussian model:
plot(lm(mass.diss.net ~ dissect.age.z + treatment + site, data=i))
# GEE model with Gaussian distribution, taking into account the non-independence of
   siblings by using the 'exchangeable correlation' structure; including age at
   dissection as numeric predictor:
gee5= geeglm(mass.diss.net ~ dissect.age.z + treatment + site, family=gaussian,
   id=sibgroup, corstr="exchangeable", data=i[!is.na(i$mass.diss.net),])
# Treatment effects:
emmeans(gee5, trt.vs.ctrl1~treatment, adjust="fdr", type="response")
# Site effects:
(e.gee5=emmeans(gee5, pairwise~site, adjust="fdr", type="response"))
# Difference between urban and woodland habitats:
contrast(e.gee5$emmeans, list(urb_nat=c(-0.5,-0.5,0.5,0.5)))
# Do sex-reversed and concordant individuals differ?
gee5.sex= geeglm(mass.diss.net ~ dissect.age.z + sex.rev.2, family=gaussian,
   id=sibgroup, corstr="exchangeable", data=i[!is.na(i$mass.diss.net) &
   !is.na(i$sex.rev.2),])
emmeans(gee5.sex, pairwise~sex.rev.2, type="response")$contrasts
# Figure S7:
par(mfrow=c(1,2), mar=c(5,5,1,1))
with(i, plot(mass.diss.net ~ dissect.age, pch=19, col=site, xlab="Age at dissection
   (days)", ylab="Body mass at dissection (mg)"))
legend("topleft", levels(i$site), pch=19, col=c(1:4))
abline(c(-28.28, 22.24), lty=2) # Coefficients for age taken from gee5
with(i, plot(mass.diss.net ~ dissect.age, pch=19, col=sex.rev.hist, xlab="Age at
   dissection (days)", ylab="Body mass at dissection (mg)"))
legend("topleft", levels(i$sex.rev.hist), pch=19, col=c(1:5))
abline(c(-28.28, 22.24), lty=2) # Coefficients for age taken from gee5
```

**# Metamorphosis**

```
# Cox's proportional hazards model with sibgroup as random factor, treatment and site
   as fixed factors; censoring individuals that did not start metamorphosis (i.e. did
   not reach Gosner stage 42):
cox2 = coxme(Surv(meta.age, censor.met) ~ treatment + site + (1|sibgroup), data=i)
# Diagnostics for proportional hazards:
zp2=cox.zph(cox2)
plot(zp2[1])
plot(zp2[2])
# Treatment effects:
emmeans(cox2, trt.vs.ctrl1~treatment, adjust="fdr", type="response")
# Site effects:
(e2=emmeans(cox2, pairwise~site, adjust="fdr", type="response"))
# Difference between urban and woodland habitats:
contrast(e2$emmeans, list(urb_nat=c(-0.5,-0.5,0.5,0.5)))
# Figure S1:
```

```
ggsurvplot(survfit(Surv(meta.age, censor.met) ~ treatment, data=i), fun="event",
   conf.int=TRUE, xlab="Time (days)", ylab="Cumulative hazard of metamorphosis ")
# Figure S3:
ggsurvplot(survfit(Surv(meta.age, censor.met) ~ site, data=i), fun="event",
   conf.int=TRUE, xlab="Time (days)", ylab="Cumulative hazard of metamorphosis")
# Do sex-reversed and concordant individuals differ?
cox2.sex = coxme(Surv(meta.age, censor.met) ~ sex.rev.2 + treatment + (1|sibgroup),
   data=i)
emmeans(cox2.sex, pairwise~sex.rev.2, type="response")$contrasts
# Figure S8:
with(i, bwplot(meta.age ~sex.rev.3|site, data=i[i$censor.met==1,], layout=c(4,1),
   ylab="Age at metamorphosis"))
```

**# Survival**

```
# Cox's proportional hazards model with sibgroup as random factor, treatment and site
   as fixed factors; censoring euthanized individuals:
cox1 = coxme(Surv(death.age, censor.death) ~ treatment + site + (1|sibgroup), data=i)
# Diagnostics for proportional hazards:
zp1=cox.zph(cox1)
plot(zp1[1])
plot(zp1[2])
# Treatment effects:
emmeans(cox1, trt.vs.ctrl1~treatment, adjust="fdr", type="response")
# Site effects:
(e1=emmeans(cox1, pairwise~site, adjust="fdr", type="response"))
# Difference between urban and woodland habitats:
contrast(e1$emmeans, list(urb_nat=c(-0.5,-0.5,0.5,0.5)))
# Figure S2:
ggsurvplot(survfit(Surv(death.age, censor.death) ~ treatment, data=i), fun="event",
   conf.int=TRUE, xlab="Time (days)", ylab="Cumulative hazard of death")
# Figure S4:
ggsurvplot(survfit(Surv(death.age, censor.death) ~ site, data=i), fun="event",
   conf.int=TRUE, xlab="Time (days)", ylab="Cumulative hazard of death")
```

**# Sex reversal**

```
# Diagnostics for binomial model:
glm1=glm(sex.rev.2 ~ treatment+site, family=binomial(logit), data=i)
glm.check=simulateResiduals(glm1, plot=T)
# Recoding the dependent variable from factor to numeric (for binomial GEE):
i$sexrev2 = as.numeric(i$sex.rev.2)-1
# GEE model with binomial distribution, taking into account the non-independence of
   siblings by using the 'exchangeable correlation' structure:
gee1=geeglm(sexrev2 ~ treatment+site, family=binomial(logit), id=sibgroup,
   corstr="exchangeable", data=i[!is.na(i$sex.rev.2),])
# Treatment effects:
emmeans(gee1, trt.vs.ctrl1~treatment, adjust="fdr", type="response")
# Site effects:
 (e.gee1=emmeans(gee1, pairwise~site, adjust="fdr", type="response"))
# Difference between urban and woodland habitats:
contrast(e.gee1$emmeans, list(urb_nat=c(-0.5,-0.5,0.5,0.5)))
```

**# Figures presented in the paper:**

**# Figure 1**

```
# Need to run: gee2, e.gee2.treat, gee3, e.gee3.treat (see above)
# Save graph into jpeg file:
jpeg("fig_cort_mass.jpg", width=10, height=15, units="cm", pointsize=10, quality=100,
   res=150)
par(mfrow=c(2,1), mar=c(0,5,0,1), oma=c(4,0,1,0), las=0)
#cort
with(i, boxplot(cort.rate~treatment, log="y", ylim=c(0.1,20), xaxt="n", yaxt="n",
   ylab="", xlab="", col="white", range=0))
par(las=1)
axis(2, at=c(0.1,1,10,100))
par(las=0)
mtext("CORT release rate (pg/mg/h)", side=2, line=3.5)
points(x=1:5, y=summary(e.gee2.treat$emmeans, level=0.84)[,2], pch=19, col="dark gray",
   cex=1.5)
arrows(1:5, summary(e.gee2.treat$emmeans, level=0.84)[,5], 1:5,
   summary(e.gee2.treat$emmeans, level=0.84)[,6], angle=90, lty=1, code=3, lwd=2,
   col="dark gray", length=0.1)
legend(legend="A", "topright", bty="n")
#post-treatment mass
with(i, boxplot(mass.may12~treatment, ylim=c(35,525), yaxt="n", ylab="", xlab="",
   names=c("0","0.01","10","100","1000"), col="white", range=0))
points(x=1:5, y=summary(e.gee3.treat$emmeans, level=0.84)[,2], pch=19, col="dark gray",
   cex=1.5)
arrows(1:5, summary(e.gee3.treat$emmeans, level=0.84)[,5], 1:5,
   summary(e.gee3.treat$emmeans, level=0.84)[,6], angle=90, lty=1, code=3, lwd=2,
   col="dark gray", length=0.1)
par(las=1)
axis(2, at=c(100,200,300,400,500))
par(las=0)
mtext("Body mass (mg)", side=2, line=3.5)
mtext("Treatment CORT (nM)", side=1, line=2.5)
legend(legend="B", "topright", bty="n")
dev.off()
```

**# Figure 2**

```
# Calculate proportions of each combination of genetic and phenotypic sex for each
   treatment:
bla=with(i, table(sex.rev.2, treatment))
bling=with(i, table(HRM.sex, treatment))
sexrevrates = bla[2,] / bling[1,]
sexrevrates*100
(sexrev.n = with(i, table(treatment, sex.rev.hist)) )
(sexrev = sexrev.n/apply(sexrev.n,1,sum))
(sexrev.n.treats = apply(sexrev.n,1,sum))
# Save graph into jpeg file:
jpeg("fig2.jpg", width=12, height=10, units="cm", pointsize=10, quality=100, res=150)
par(mfrow=c(1,1), mar=c(5,5,3,9), las=0)
sexbarplot = with(i, barplot(t(sexrev), names.arg= c("0","0.01","10","100","1000"),
    xlab="Treatment CORT (nM)", yaxt="n", ylab="Proportion of individuals",
    cex.lab=1.2, cex.axis=1.2, cex.names=1.2, width=sexrev.n.treats, border="black",
```

```
    space=0.15, col=c(1:5), legend.text=levels(i$sex.rev.hist), args.legend =
    list(bg="white", x=380, title="Sex", title.adj=0.1, border="black", bty="n",
    cex=1)))
par(las=1)
axis(2, at=c(0,0.25,0.5,0.75,1))
par(las=0)
#abline(h=0.5, lty=2)
mtext(c("11.1%", "21.7%", "31.8%", "11.1%", "20.0%"), at=sexbarplot, side=3, line=1.5,
   cex=0.8)
mtext(c("(41)", "(38)", "(42)", "(41)", "(41)"), at=sexbarplot, side=3, line=0.5,
   cex=0.8)
dev.off()
```

**# Figure 3**

```
# Need to run: gee2, e.gee2, gee4, e.gee4 (see above)
# Calculate mortality and sex-reversal rates for each site:
y=summary(e.gee4$emmeans)[,2]
x=summary(e.gee2$emmeans)[,2]
(weights = with(i, table(site, sex.rev.2)))
(deaths = with(i, table(site, censor.death)))
(mortalities = deaths[,2] / 75)
# Save graph into jpeg file:
jpeg("fig3.jpg", width=12, height=12, units="cm", pointsize=10, quality=100, res=150)
par(mfrow=c(1,1), mar=c(5,5,1,1), las=0)
plot(c(0.4,2.1), c(360,530), type="n", yaxt="n", xlab="CORT release rate (pg/mg/h)",
   ylab="Body mass (mg)", cex.lab=1.2)
arrows(x, summary(e.gee4$emmeans, level=0.84)[,5], x, summary(e.gee4$emmeans,
   level=0.84)[,6], angle=90, lty=1, code=3, lwd=1.5, col=1, length=0.03)
arrows(summary(e.gee2$emmeans, level=0.84)[,5], y, summary(e.gee2$emmeans,
   level=0.84)[,6], y, angle=90, lty=1, code=3, lwd=1.5, col=1, length=0.03)
for (j in 1:length(x)) {
floating.pie(xpos=x[j], ypos=y[j], x=weights[j,], col=c(gray(1-mortalities[j]),
   "black"), radius=0.08)
}
text(c("I: 2.4%", "B: 23.3%", "G: 30.8%", "E: 33.3%"), x=rep(x+0.02,4),
   y=rep(summary(e.gee4$emmeans, level=0.84)[,6] + 2, 4), cex=1, pos=3)
par(las=1)
axis(2, at=c(400,450,500))
par(las=0)
text("Woodland habitats: I, B", x=1.3, y=525, pos=4)
text("Urban habitats: G, E", x=1.3, y=515, pos=4)
dev.off()
```
