## Supplementary table & figures, data, and R script. for "Does stress make males? An experiment on the role of glucocorticoids in anuran sex reversal": Bokony_Supplementary CORTsex.pdf

**Table S1.** Differences between sex-concordant individuals (XY males and XX females) and sex-reversed individuals (XX genotype with testes or ovotestes), estimated as linear contrasts of model-estimated marginal means.

| Dependent variable | Contrast ± SE | P |
|---|---|---|
| CORT release rate* | 0.7 ± 0.22 | 0.261 |
| Time to metamorphosis‡ | 1.3 ± 0.40 | 0.384 |
| Body mass at the end of treatment† | 2.8 ± 20.7 | 0.890 |
| Body mass at metamorphosis† | 14.3 ± 9.43 | 0.130 |
| Body mass at dissection† | 52.4 ± 39.9 | 0.188 |

* Ratios of group means are given from Generalized Estimating Equations model (concordant/sex-reversed).

‡ Hazard ratios are given from Cox's proportional hazards model (concordant/sex-reversed).

† Differences between group means are given from Generalized Estimating Equations models (concordant – sex-reversed).

**Figure S1.** Kaplan-Meier curves of time to metamorphosis in the five corticosterone treatment groups, with 95% confidence intervals.

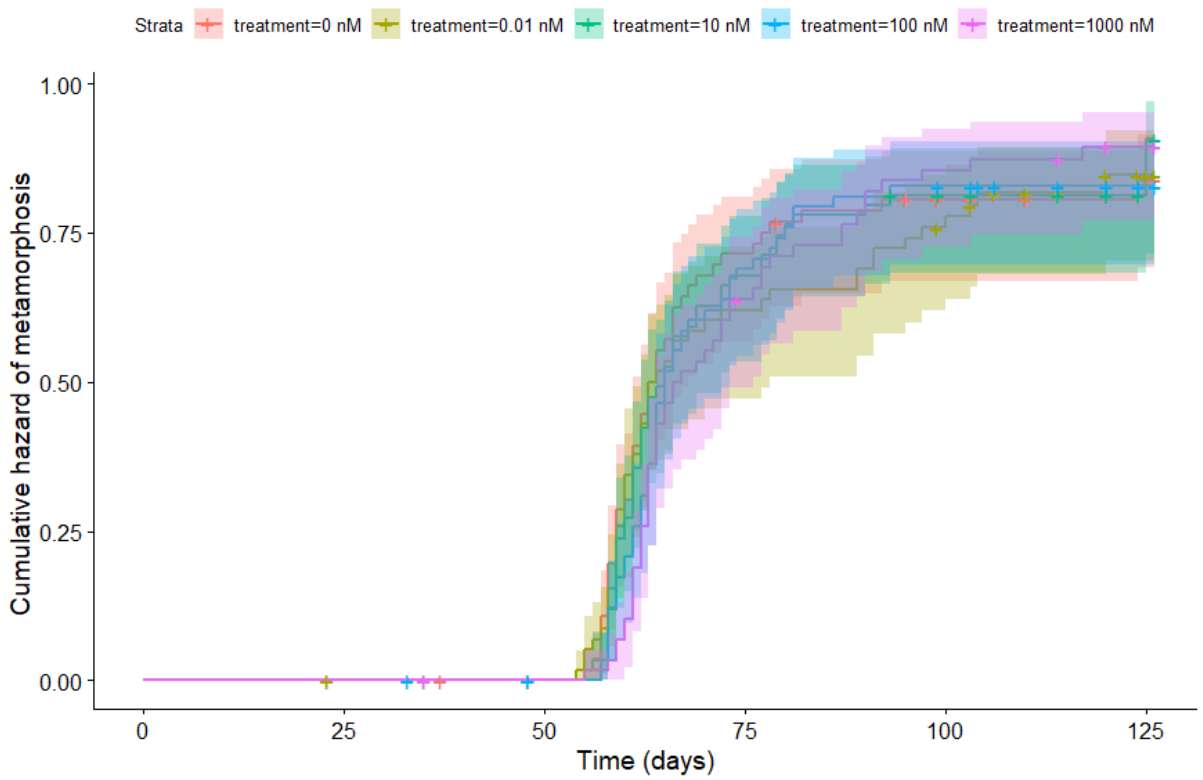


**Figure S2.** Kaplan-Meier curves of survival over the experiment in the five corticosterone treatment groups, with 95% confidence intervals.

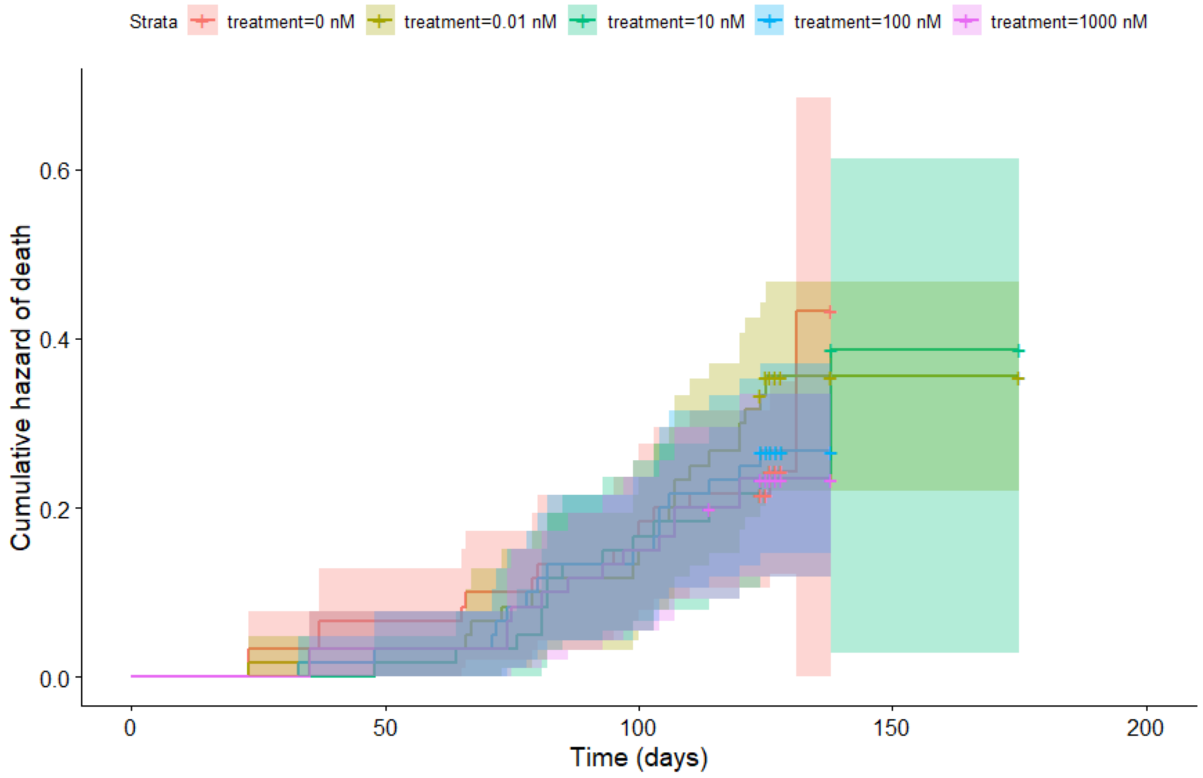

**Figure S3.** Kaplan-Meier curves of time to metamorphosis by the site of origin, with 95% confidence intervals.

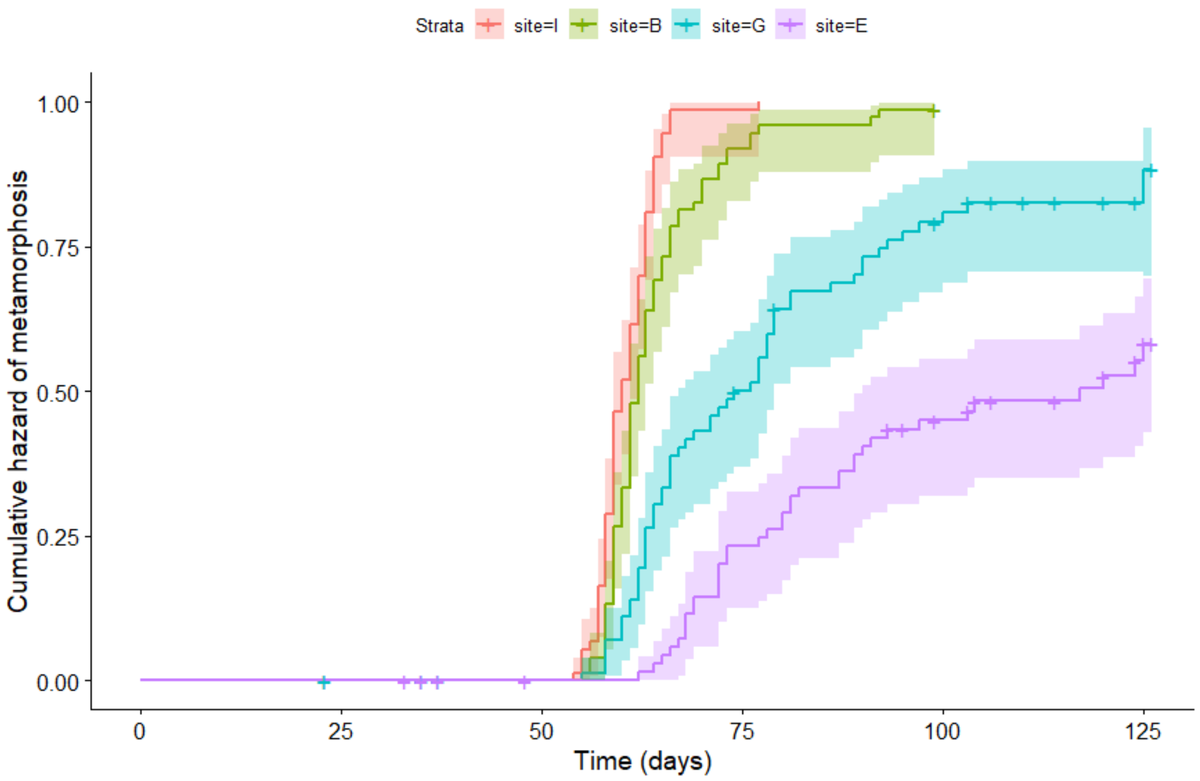


**Figure S4.** Kaplan-Meier curves of survival over the experiment by the site of origin, with 95% confidence intervals.

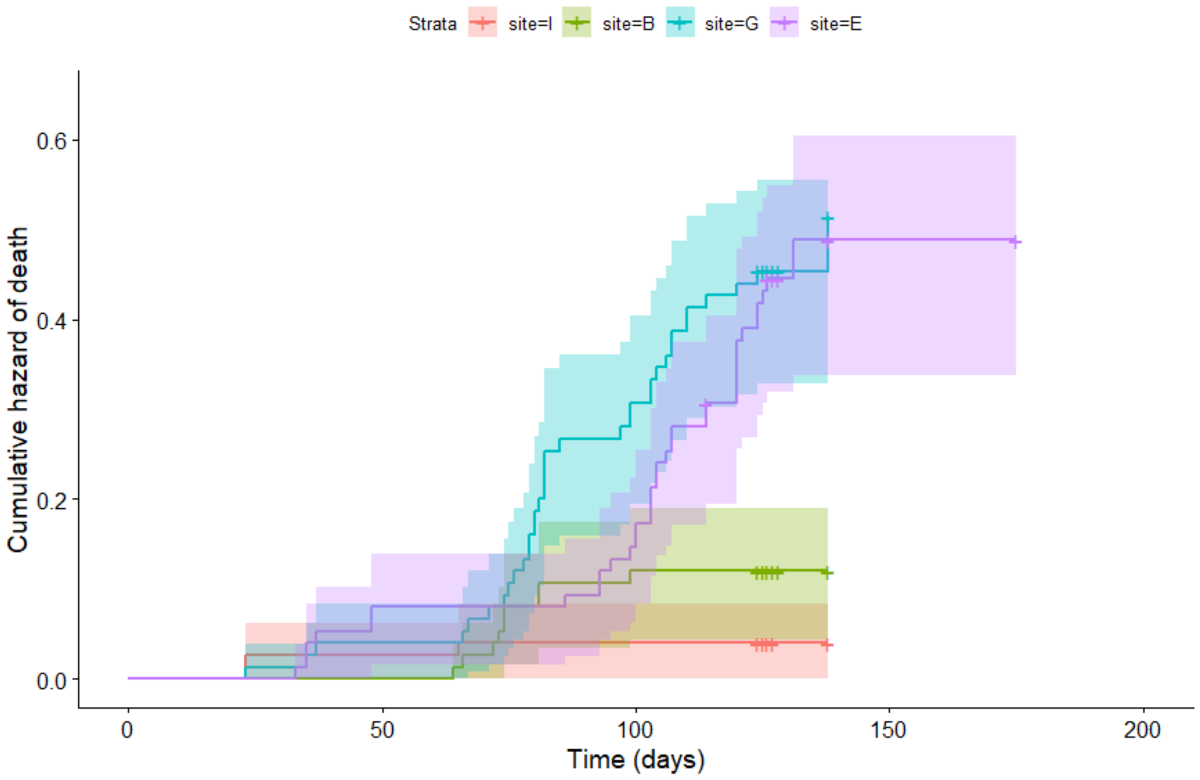

**Figure S5.** Tadpole corticosterone release rates at the end of treatment, by site of origin and combination of genetic and phenotypic sex. In each boxplot, the thick horizontal line, box, and whiskers represent the median, interquartile range, and data range, respectively. The category „XX male" includes XX individuals with testes or ovotestes.

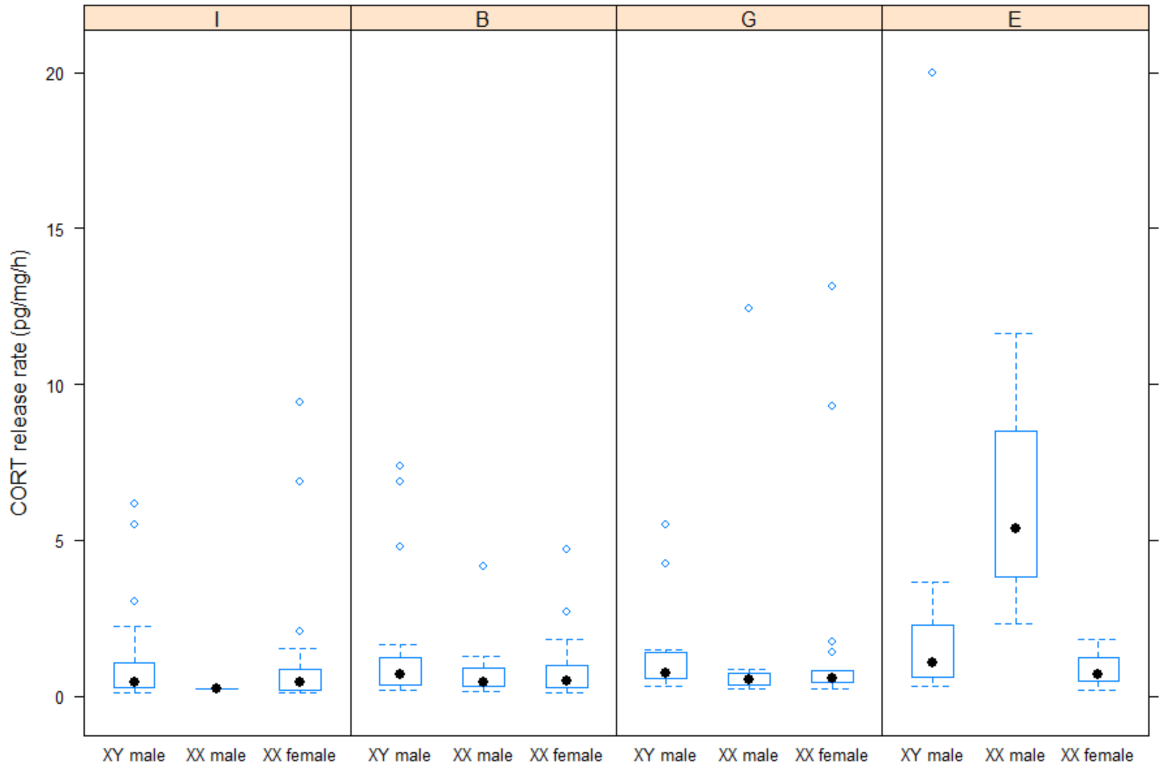

**Figure S6.** Body mass at the end of treatment, upon metamorphosis, and at dissection, by site of origin and combination of genetic and phenotypic sex. In each boxplot, the thick horizontal line, box, and whiskers represent the median, interquartile range, and data range, respectively. The category „XX male” includes XX individuals with testes or ovotestes.

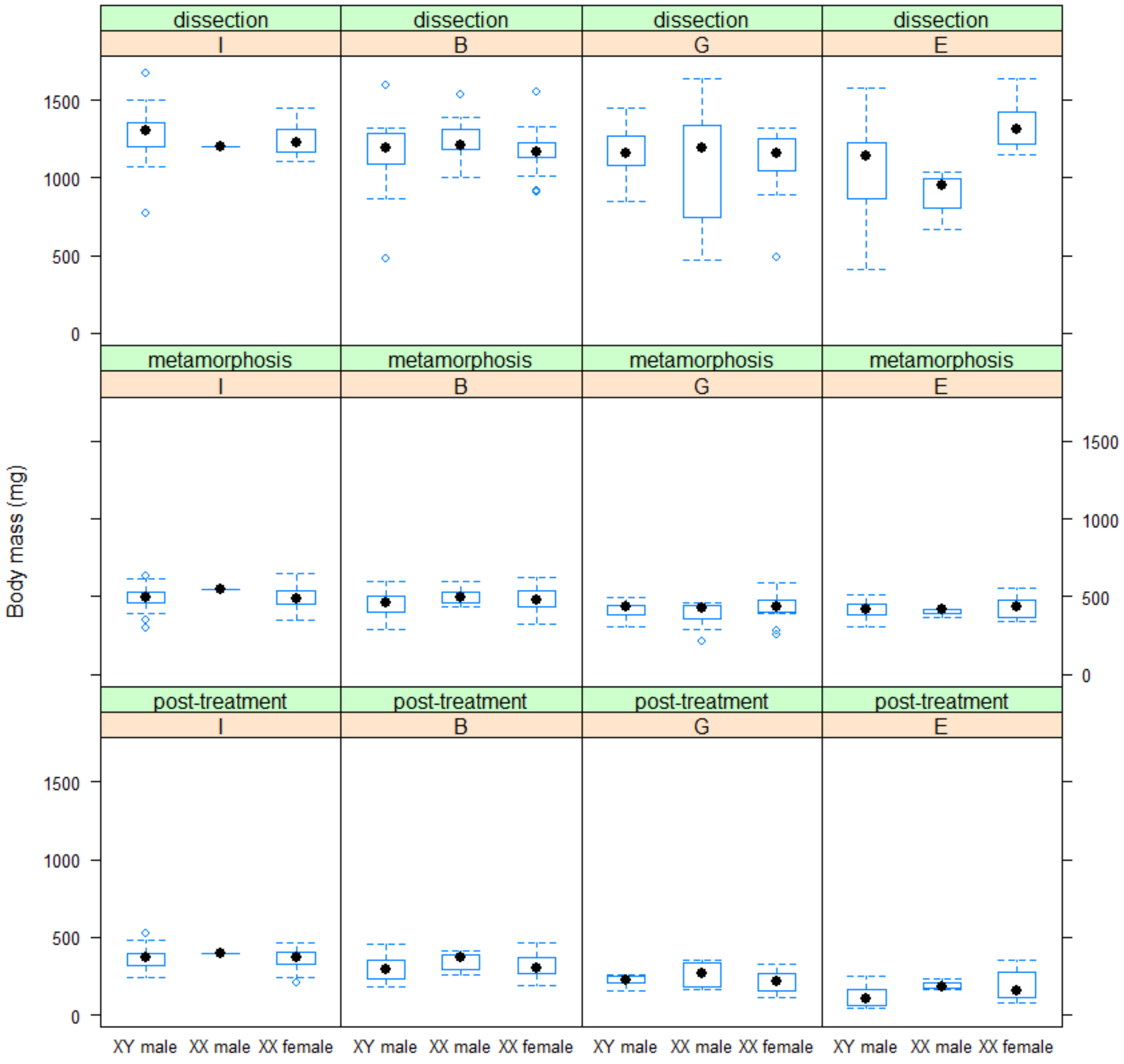

**Figure S7.** Body mass at at dissection as a function of age at dissection, by site of origin (on the left) and by combination of genetic and phenotypic sex (on the right). The dashed lines were fitted by a Generalized Estimating Equations model (including treatment, site, and age of dissection as predictors).

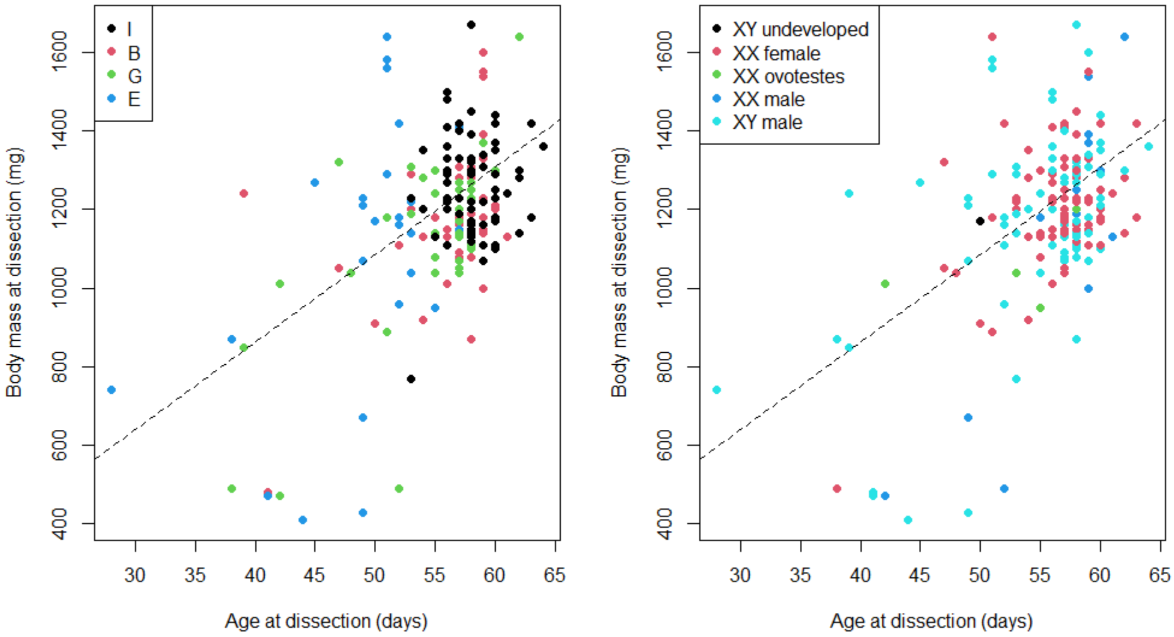

**Figure S8.** Age at metamorphosis (number of days from Gosner stage 25 to Gosner stage 42) by site of origin and combination of genetic and phenotypic sex. In each boxplot, the thick horizontal line, box, and whiskers represent the median, interquartile range, and data range, respectively. Individuals that did not metamorphose are not shown here. The category „XX male" includes XX individuals with testes or ovotestes.

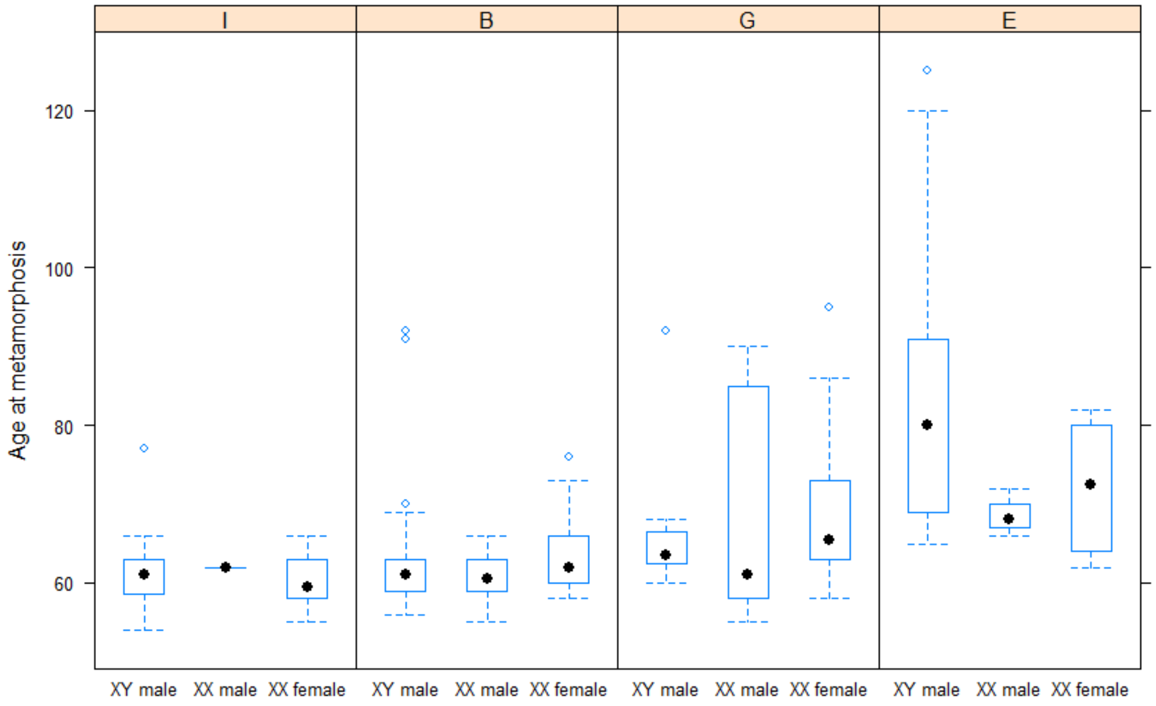
